# Relationships between Human Brain Structural Connectomes and Traits

**DOI:** 10.1101/256933

**Authors:** Zhengwu Zhang, Genevera I. Allen, Hongtu Zhu, David Dunson

**Author notes:** Correspondence: Dr. Zhengwu Zhang, University of Rochester, Department of Biostatistics & Computational Biology, Rochester, NY, USA.

## Abstract

Advanced brain imaging techniques make it possible to measure individuals’ structural connectomes in large cohort studies non-invasively. However, due to limitations in image resolution and pre-processing, questions remain about whether reconstructed connectomes are measured accurately enough to detect relationships with human traits and behaviors. Using a state-of-the-art structural connectome processing pipeline and a novel dimensionality reduction technique applied to data from the Human Connectome Project (HCP), we show strong relationships between connectome structure and various human traits. Our dimensionality reduction approach uses a tensor characterization of the connectomes and relies on a generalization of principal components analysis. We analyze over 1100 scans for 1076 subjects from the HCP and the Sherbrooke test-retest data set as well as 175 human traits that measure domains including cognition, substance use, motor, sensory and emotion. We find that brain connectomes are associated with many traits. Specifically, fluid intelligence, language comprehension, and motor skills are associated with increased cortical-cortical brain connectivity, while the use of alcohol, tobacco, and marijuana are associated with decreased cortical-cortical connectivity.

## Introduction

The human brain structural *connectome*, defined here as the collection of white matter fiber tracts connecting different regions of the brain ^[1-4]^, plays a crucial role in how the brain responds to everyday tasks and life’s challenges. There has been a huge interest in studying connectomes and understanding how they vary for individuals in different groups according to traits and substance exposures. Such studies have typically focused on functional connectomes instead of structural connectomes ^[1,5-8]^, due to the difficulty of recovering reliable structural connectomes ^[9,10]^. Recent advances in noninvasive brain imaging and preprocessing have produced sophisticated tools to routinely measure brain structural connectomes for different individuals. Relying on high quality imaging data and many different traits for a large number of study participants obtained in the Human Connectome Project (HCP) ^[11,12]^, this article focuses on analyzing relationships between brain structural connectomes and different traits and substance exposures using novel data science tools applied to the HCP data.

Estimation of the structural connectome relies on a combination of diffusion magnetic resonance imaging (dMRI) and structural MRI (sMRI). dMRI collects information on the directional diffusion of water molecules in the brain ^[13-15]^. As diffusion tends to occur in a directional manner along fiber tracts in white matter, while being essentially non-directional in gray matter, dMRI provides information on locations of fiber tracts.

Tractography ^[16-18]^ extracts the tracts from dMRI data, yielding a very large number of ‘tubes’ snaking through the brain. These data are enormous and complex, and it is difficult to align such data for different subjects into a common coordinate system, which is necessary for statistical analysis. For these reasons, it is typical to parcellate the brain into anatomical regions of interest (ROIs) using sMRI ^[19,20]^ according to pre-defined templates ^[19-21]^. This allows alignment of different individuals, and connectome data reduction into a connectivity matrix.

Advances in imaging techniques ^[22]^ and preprocessing pipelines ^[18,23-25]^ have improved the reconstruction of structural connectomes ^[25]^. However, reconstruction errors inevitably occur ^[9,10,26]^ and questions remain about whether existing reconstruction and analysis methods are sufficient to detect (potentially subtle) associations between connectomes and human traits. Current statistical approaches focus primarily on reducing the connectome to a binary adjacency matrix containing 0-1 indicators of any fiber connections between ROIs. The adjacency matrix is then further reduced to topological summary statistics of the brain graph, providing low-dimensional numerical summaries to be used in statistical analyses ^[27-29]^. Such connectome simplification is appealing due to its interpretability, but leads to an enormous loss of information.

In our framework of analyzing brain connectomes and human traits, we rely on a new structural connectome processing pipeline, and improved methods for representing brain connectomes ^[24]^. In particular, we use a *tensor network* representation that incorporates multiple features measuring the strength and nature of white matter tracts between each pair of brain ROIs. This representation better preserves information in the tractography data, and allows flexibility in examining associations with traits. We extend principal components analysis (PCA) to tensor network data via a semi-symmetric tensor decomposition method, which produces brain connectome PC scores for each subject. These scores can be used for visualization and efficient inference on relationships between connectomes and human traits.

Based on our analysis of data on 1076 individuals and 175 traits, we find a strong relationship between structural cortical-cortical connectomes and multiple traits, particularly those related to cognition, motion and substance usage. Extensive predictive analyses also confirm that the structural connectomes can significantly improve the prediction of these traits in addition to demographic predictors such as age and gender. Further investigation shows that traits related to positive lifestyles, such as good reading ability, high fluid intelligence, and good motor skills, tend to have positive correlations on cortical-cortical brain connections. On the other hand, substance use, including binge drinking, tobacco, and marijuana use, can reduce interconnections. These findings are consistent with related analyses for functional connectomes ^[5,7]^.

## Results

### Data description

From the raw diffusion MRI (dMRI) and structural MRI (sMRI) data, we want to reliably extract the white matter fiber tracts representing an individual’s brain structural connectome. Figure 1 illustrates our data processing pipeline. The first step is to use a reproducible probabilistic tractography algorithm ^[9,18]^ to generate fiber tracts (also called streamlines) across the whole brain using anatomical information to reduce bias ^[18]^. The well-known Desikan-Killiany atlas ^[19]^ is then used to define brain ROIs relying on Freesurfer software, leading to 68 cortical surface ROIs with 34 in each hemisphere. Figure 1 (a) illustrates the Desikan-Killiany parcellation and a reconstructed connectome.

For each pair of ROIs, we identify the streamlines connecting them. Several procedures are used to increase the robustness and reproducibility of the extracted streamlines for each ROI pair: (1) each gray matter ROI is dilated to include a small portion of the white matter region ^[10,24]^; (2) streamlines connecting multiple ROIs are cut into pieces such that we can extract the complete pathway between any ROI pair and (3) apparent outlier streamlines are removed. Extensive experiments have illustrated that these procedures can significantly improve the robustness of extracted connectomes and their reproducibility ^[24]^.

**Figure 1:**
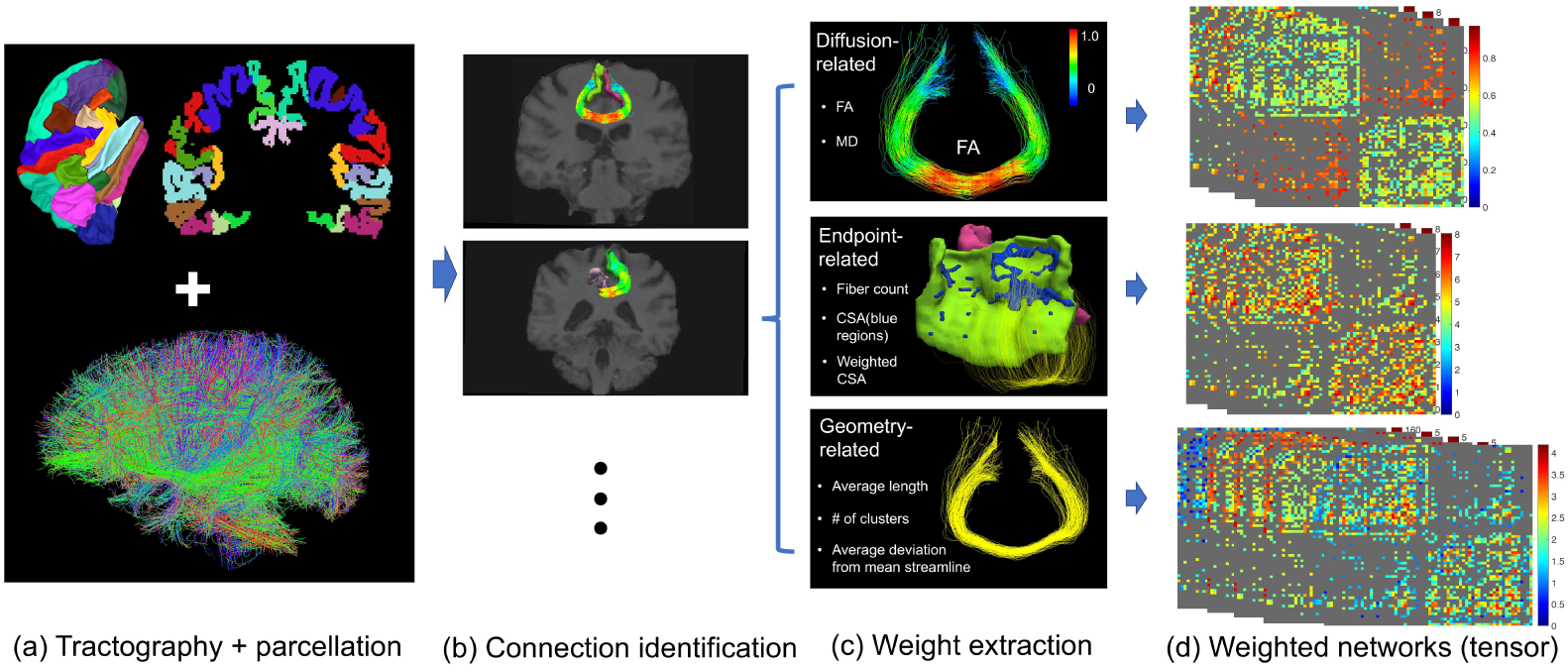
Pipeline of the preprocessing steps to extract weighted networks from the dMRI and sMRI image data. (a) Desikan-Killiany parcellation and the tractography data for an individual’s brain; (b) extraction of streamlines between two ROIs; (c) feature extraction from each connection and (d) extracted weighted networks.

For a particular pair of ROIs, instead of only using the streamline count as the connection weight, we extract multiple features from the streamlines to describe the connection. These features can be classified into three groups: (i) endpoint-related features, (ii) diffusion-related features and (iii) geometry-related features (as illustrated in Figure 1 (c)). The online method section introduces more details about these features. For each individual and feature, one can obtain a weighted adjacency matrix representation of the brain connectome. If we stack together the matrices corresponding to different features on the same individual’s brain, we obtain a three-way semi-symmetric tensor network with a dimensionality of *P* × *P* × *M*, where *P* is the number of ROIs (nodes in the brain network) and *M* is the number of features describing connections between each pair of nodes.

Applying this pipeline, we processed the Sherbrooke test-retest data set with 11 subjects and 3 repeated scans per subject (for our reproducibility study), and the HCP data set with 1065 subjects (from the latest release of HCP data set in 2017) ^[30]^. For each subject, 12 weighted networks are extracted: three endpoint-related features (count of streamlines, connected surface area (CSA) and weighted CSA), four diffusion-related features (mean and max values of fractional anisotropy (FA) and mean diffusivity (MD)), and five geometry-related features (cluster number, average length and mean deviations from a template streamline). In addition, 175 trait measures for each subject in the HCP data set were also obtained. The 175 trait measures are from eight categories describing a human’s status and behavior: cognition, motor, substance use, psychiatric and life function, sense, emotion, personality, and health. A detailed description of these traits is included in the Excel spreadsheet of Supplementary Material II. In total, we extract 13, 176 networks and 186, 375 trait scores for our statistical analyses.

### Tensor network principal components analysis

Our focus is on inferring relationships between brain structural connectomes and human traits. To address this goal using the extracted connectome representation, it is necessary to develop a statistical approach to assessing associations between tensor network representations of the brain connectome and traits. A key problem in this respect is how to estimate low-dimensional features summarizing the brain tensor network without loosing valuable information. If the tensor network for each individual could be reorganized into a vector, Principal Components Analysis (PCA) could be used to extract brain connectome PC scores for each individual. However, vectorizing the tensor would discard valuable information about the network structure. Instead, we propose to use a semi-symmetric tensor generalization of PCA (see the online method section for more details).

Our Tensor Network PCA (TN-PCA) approach approximates the brain tensor network using *K* components, with the components ordered to have decreasing impact similarly to common practice in PCA. Individuals are assigned a brain connectome PC score for each of the *K* components, measuring the extent to which their brain tensor network expresses that particular tensor network (TN) component. Each TN component is itself a tensor network, but one having a very simple rank one structure depending on scores for each brain ROI. By using TN-PCA, we can replace the high dimensional tensor network summarizing an individual’s brain connectome with a *K*-dimensional vector of brain PC scores; these scores can be used for visualizing variation among individuals in their brain connectomes and in statistical analyses studying relationships between connectomes and traits.

Taking one feature, CSA, as an example (M, the third dimension of tensor network representation, degenerates to one), Figure 2 (a) shows brain PC scores for each of the 33 CSA networks in the Sherbrooke test-retest data set using *K* = 3 for visualization. Each combination of color and marker type represents three scans from the same subject (11 unique ones for 11 subjects). Even using only *K* = 3 components, the CSA brain networks display a clear clustering pattern, suggesting that not only are the extracted connectomes from the repeated scans reproducible but also that we can distinguish between different subjects based on only three components. To formally assess this, we applied nearest neighbor clustering to the PC scores for the 12 types of weighted networks separately (M=1) and jointly (M=12) under different *K*. Figure 2 (b) shows the results. With a moderate *K*, e.g, *K* = 10, we can almost perfectly cluster the repeated scans of all 11 subjects based on the discriminative features such as CSA, count and cluster number. The ranking in the classification results for *K* = 10 gives us an idea of the discriminative power of each feature: CSA > Weighted CSA > Count > Cluster No. > others.

**Figure 2:**
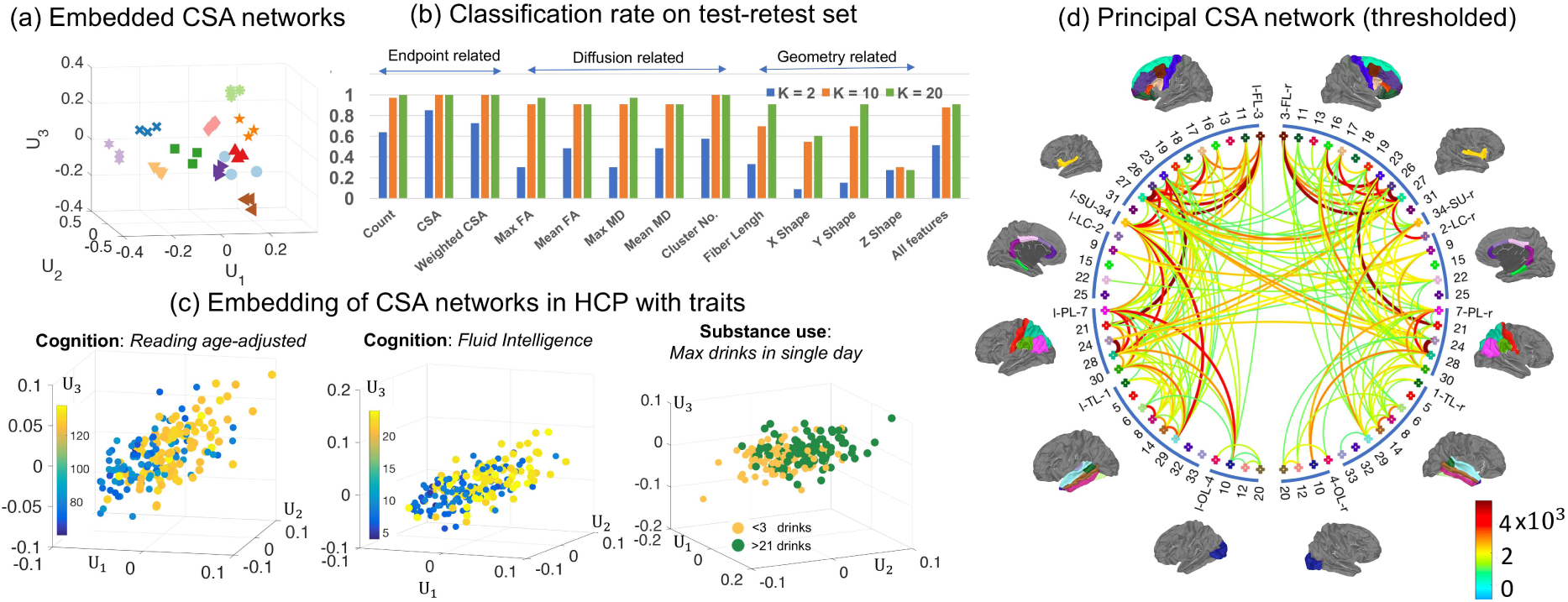
Tensor Network (TN-) Principal Components Analysis (PCA). In (a), we display brain PC scores for all 33 CSA networks from the Sherbrooke test-retest data set. Each unique combination of color and marker type represents three scans from the same subject (11 unique ones). In (b), we show nearest neighbor clustering results using the brain PC scores for all 12 types of weighted networks separately and jointly under different *K*. In (c), we show PC scores with traits. For each cognition trait, we selected 100 subjects with low trait scores and 100 subjects with high scores. For the substance use trait (alcohol use), we selected subjects with < 3 drinks and subjects with > 21 drinks. In (d), we display the principal CSA brain network calculated using all 1065 subjects from the HCP data set. For display purposes, we thresholded the dense principal brain network to keep only the 200 most connected edges.

TN-PCA allows one to visualize relationships between the structural connectome and various traits in the HCP data. Taking the CSA network as one example, Figure 2 (c) displays the first three brain PC scores along with three selected traits (two cognition, one substance use). For the two cognition traits (oral reading test score and fluid intelligence), we selected 100 subjects with low trait scores and 100 subjects with high scores and plotted their brain PC scores. For the substance use trait (max drinks in a single day), we plot subjects with low (< 3 alcohol drinks) and high values (> 21 alcohol drinks). We can clearly observe separation between different groups of subjects in these plots, indicating that brain connection patterns are different for these two groups (measured by the CSA feature; we have similar findings on some other features, e.g. the fiber count).

Principal brain networks can also be obtained as a byproduct of TN-PCA. We define the rank *K* principal brain network to be the network given by the sum of the first *K* rank-one tensor network components from TN-PCA (see the online method section for more details). Similar to examining patterns amongst features by exploring the PC loadings, the principal brain network exhibits major patterns of structural connectivity that explain most of the variation across the population. Figure 2 (d) displays the thresholded principal brain network derived from the CSA networks of 1065 subjects from the HCP data set; here, we take *K* = 30 and threshold the edges so that the 200 most connected pairs of brain regions are displayed. The principal brain network gives a visual summary of all the structural connectomes in the HCP population. Compared to the mean network for the HCP population (shown in Figure 3 in the supplemental materials), our principal brain network yields additional insights into which structural connections tend to have major differences across subjects. Specifically, we see that there is large variation in the strength of connections within hemispheres as well as a few strong inter-hemispheric connections that vary across subjects. As we will investigate in the next sections, these variations in brain connections across subjects may be related to differences in traits.

### Relating brain connectomes with traits

To study relationships between brain connectomes and traits, we use the extracted brain PC score vector for each subject and feature. We focus on the top 30 PC scores, which explain approximately 90% of the variation in the brain tensor network (for most of the features). We first assess whether there are significant differences in the distribution of the brain PC scores among subjects having a low versus high value for each trait. Out of the 1065 HCP subjects, we identified groups of 100 having the highest and lowest values for each trait. We used the Maximum Mean Discrepancy (MMD) test ^[31]^ to obtain *p* values for differences in the brain connectomes across these two groups for each trait. Results are shown in Figure 3. Figure 3 (a) shows *p* values for the CSA weighted networks. Different thresholds for significance based on false discovery rate (FDR) control using ^[32]^ are marked with different colored lines. Corresponding results for 7 types of weighted networks (using endpoint and diffusion related features) are shown in Figure 3 (b). We exclude the remaining 5 geometry-related features because of their relatively poor performance in our analyses of the test-retest data (refer to Figure 2).

Based on these results, many traits are significantly related to brain structural connectomes. Traits in the same domain are placed next to each other in Figure 3 (b). The block patterns of significance add evidence that brain structural connectomes relate more broadly to these trait domains. In the cognitive domain, structural connectomes are related to fluid intelligence, language decoding and comprehension, working memory, and some executive functions. In the motor domain, connectomes are related to endurance and strength. In the substance use domain, connectomes are related to alcohol consumption, tobacco, illicit drug and marijuana use. In the sensory domain, connectomes are related to hearing and taste. In the emotion domain, connectomes are related to negative emotions, such as anger and anxiety. In the health domain, connectomes are related to height, weight and BMI. It is clear that endpoint-related features are more discriminative according to the hypothesis testing results; more traits are significant adjusting for false discoveries using features such as count, CSA and weighted CSA than when using diffusion-related features.

Having established significant relationships between brain connectomes and traits, it is important to assess how well traits can be predicted based on connectomes. A baseline model that only uses demographic variables of age and gender as predictors is compared with a full model that also includes brain connectome PC scores. The 1065 subjects in the HCP data set are randomly divided into three groups: a training group containing 66% of the subjects, a validation group containing 17%, and a test group containing 17%. We trained various machine learning algorithms for prediction using the training data set, with the number of components *K* treated as a tuning parameter. The prediction improvement of the full model over the baseline model is measured using the relative ratio *ρ* (more details on calculating *ρ* are presented in the online method section). For each trait, the best model (with the highest *ρ*) is selected based on the validation data set. Panel (a) of Supplementary Figure 4 presents the results (*ρ*’s for different traits and features) for the validation and test data sets based on the average of 50 runs.

**Figure 3:**
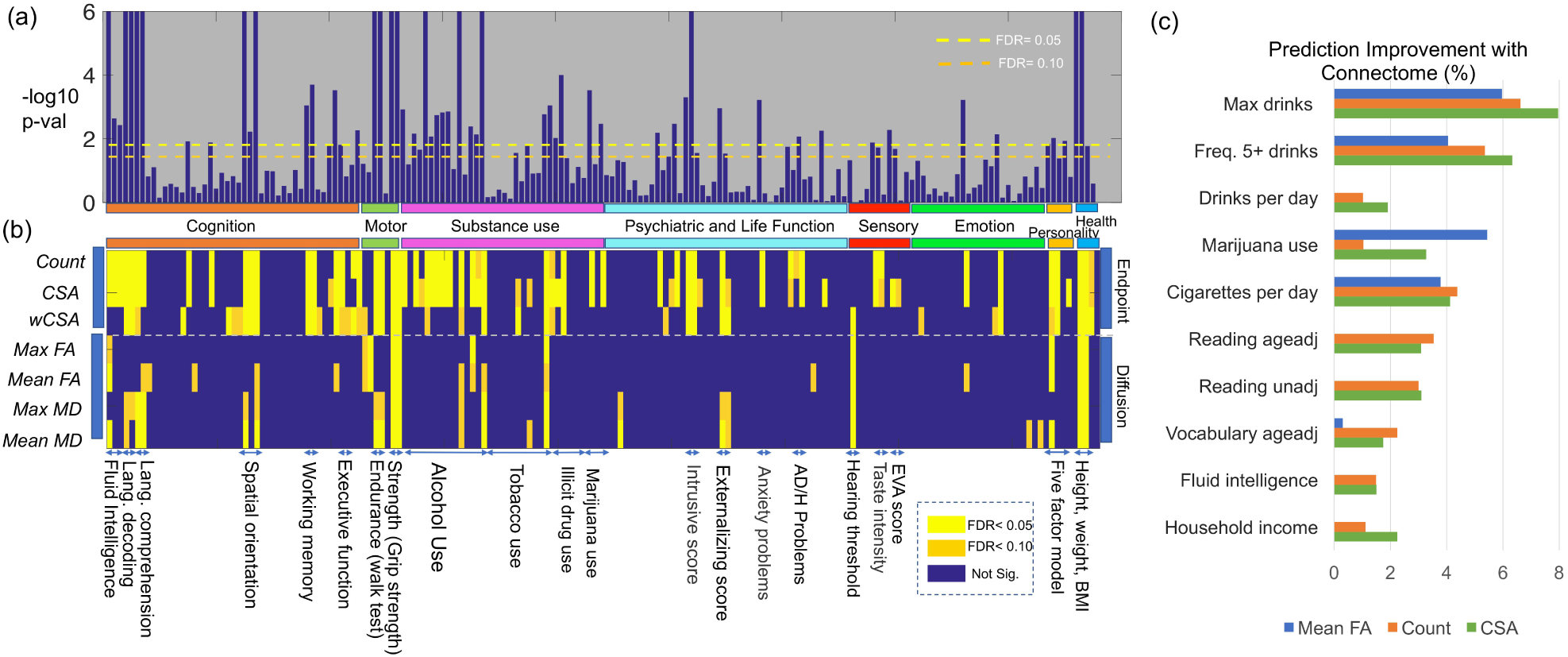
Relation between structural brain connectomes and various traits. Panels (a) and (b) show hypothesis testing results for 175 traits and 7 different weighted networks. In (a), we show p-values (–log_10_ scale) of the CSA weighted networks with different traits. Two different FDR thresholds are used (different colored dash lines). In (b), we present hypothesis testing results for different combinations of traits and networks. Each row shows a type of weighted network and each column shows a particular trait. The significance is displayed based on different FDR values. Panel (c) shows the top 10 traits in terms of their predictability based on brain connectomes adjusting for age and gender as covariates. The prediction improvement ratio *ρ*_*p*_ for the *p*th trait is shown.

Structural brain connections improve the prediction of many traits beyond that of just age and gender; these results align with those in Figure 3. According to the values of *ρ* in the test data set, we selected the 10 traits yielding the largest predictive improvements based on connectomes. The *ρ*’s for these 10 traits are displayed in Figure 3 panel (c). The 10 traits come from two domains: substance use (5 traits) and cognition (5 traits). Among the five traits of substance use, three of them are related to alcohol use, one to cigarette use and the last one to marijuana use. A close inspection of the two alcohol use traits was performed and the results are presented in the Supplementary Figure 4 panels (c) and (d). Consider the trait that measures lifetime max drinks in a single day as one example; it is an ordinal variable ranging from 1 to 7, with 1 corresponding to less than 3 drinks and 7 to more than 21 drinks. For subjects with reported values from 1 to 7, our model based on the brain connectome PC scores predicted means values (on the test data set) for each group of 1.97, 1.83, 2.27, 2.67, 2.91, 3.13, 4.04, respectively, showing a clear increasing pattern. In another example presented in Figure 4, we extract subjects with value 1 (light drinkers, totalling 191) and 7 (binge drinkers, totalling 93) and use LDA to perform binary classification. Based on the brain connectome PC scores alone, we obtain a classification accuracy of 80.99%. These results clearly indicate that using only the brain connectomes, we can distinguish with surprising accuracy between individuals with low and binge alcohol consumption.

Of the five leading cognitive traits, three are related to language and vocabulary decoding ability, one to fluid intelligence, and the other to household income (we consider the household income as a trait that is loosely connected with cognition). A close inspection of the language decoding trait is presented in the Supplementary Figure 4 panel (b). The language decoding trait score (after age adjustment) can be predicted ~4% better (*p* < 0.0002) under a random forest model with the additional CSA brain PC scores (the random forest model is selected based on validation data). On the test data set, the correlation between the predicted trait and the subject self-reported trait is *r* = 0.27 (based on the average of ten runs). If we restrict the analysis to the 200 subjects with the highest and lowest traits (plotted in the first column of panel (c) in Figure 2), the correlation increases to *r* = 0.45. Similar results are observed for the traits of fluid intelligence and vocabulary decoding. Given that these trait scores are only a rough measure of a person’s cognitive function, the results are very promising, indicating that brain structural connectomes can partially account differences in cognitive abilities.

### Interpreting relationships between traits and connectomes

Having established that a particular trait is significantly associated with the brain connectome, it is important to infer how the connectome varies across levels of the trait. For example, is the association specific to certain sub-networks and in what direction is the association? We start by studying how brain connectome PC scores vary with the trait, and then map these changes back onto the network. The approach taken depends on the measurement scale of the trait, which is continuous, ordinal or binary (see the Supplementary Material II for more details).

For continuous variables we use canonical correlation analysis (CCA) ^[33]^, while for categorical variables we rely on linear discriminant analysis (LDA), to infer the direction of change in brain PC scores with increasing values of the trait. Letting U_*K*_(*i*,:) denote the *K* PC scores for individual *i*, we estimate a unit direction w such that 〈w, U_*K*_(*i*,:)〉 has maximal correlation with the trait for CCA and maximal separation between groups for LDA. To map this direction w back on to the brain network for interpretability, we let 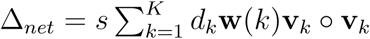, where *d*_*k*_ and v_*k*_ come from the TN-PCA analysis, and *s* is a scaling parameter (the online method section presents more details on calculating *s*). ∆_*net*_ describes how the network changes with increases of a particular trait.

The following analyses are based on *K* = 30 (results are robust for *K* ~ 20 – 60). We selected three representative traits among the 10 most predictable traits based on connectomes (see above): language reading age adjusted score, lifetime max drinks consumed in a single day and the use of marijuana. The language reading score is a continuous variable, and the other two are categorical. We study how the CSA network changes with increases in these traits.

For the language reading trait, the plot of the top 3 PC scores for the 100 subjects with the highest scores and 100 with the lowest scores is shown in panel (c) of Figure 2. Panel (a) in Figure 4 shows the network change (∆_*net*_) with the trait; we show only the 50 edges that change most. There are four major white matter connections that increase significantly with reading ability. They connect the left and right frontal lobes (FL) and parietal lobes (PL): (rFL, lFL), (rFL, rPL), (lFL, lPL), (rPL, lPL). On closer inspection, the strongest connections are among left and right brain nodes of 27-superior frontal, 26-middle frontal (BA 46), 28-superior parietal and 24-precuneus ( Supplementary Material III has detailed information on each ROI). These regions have shown strong associations with language ability in previous studies of Positron Emission Tomography (PET), functional MRI ^[8,34]^ and cortical thickness ^[35]^. There were no edges that decreased significantly with increasing reading ability. This result is robust to adjustment for age and gender.

Lifetime max drinks is an ordinal variable ranging from 1 to 7, with 1 corresponding to less than 3 drinks and 7 more than 21 drinks. The third column in Figure 2 (c) shows the top 3 brain PC scores for individuals with a max drink score of 1 (191 light drinkers) and 7 (93 binge drinkers). The separation between light and binge drinkers suggests structural connectome differences between the two groups. Panel (b) in Figure 4 shows the result of using LDA to identify brain network changes between light and binge drinkers (∆_*net*_). As max drinks increases, inter-hemisphere connections (especially the connections between rFL, lFL, rPL, and lPL) decrease. Different from the language reading trait, we observe that ∆_*net*_ for the alcohol drinking are mostly associated with the frontal lobes (almost all connections are involved with either rFL and lFL). Previous studies ^[36]^ have provided evidence of relationships between alcoholism and dysfunction and deficits in the frontal lobe. We repeated this analysis for marijuana use (583 used and 481 did not), and the result is shown in panel (c) of Figure 4. Similarly to alcohol, marijuana use is also associated with decreases in inter-hemisphere connections.

To assess how well subjects with different trait scores can be distinguished based on their brain connectomes, we calculated the correlation between traits and their predictive values for continuous traits and the classification rate for categorical traits, based on 〈w, U_*K*_(*i*,:)〉. The results are presented in Figure 4. The correlation between the projected U_*K*_(*i*,:) and the language reading score is 0.45 for the 200 subjects, indicating a strong relation between reading ability and the subjects’ brain networks. The classification rate is 80.99% for binge versus light drinking. More specifically, the sensitivity (binge drinker is identified as binge drinker) is 59.1% and the specificity (light drinker is identified as light drinker) is 91.6%. The classification rate for marijuana use or not is 59.68% (sensitivity: 26.8%, specificity: 86.8%) This rate is surprisingly high for alcohol, indicating sizable differences in these subjects’ brain structural connectomes, while the rate for marijuana is only slightly better than chance.

**Figure 4:**
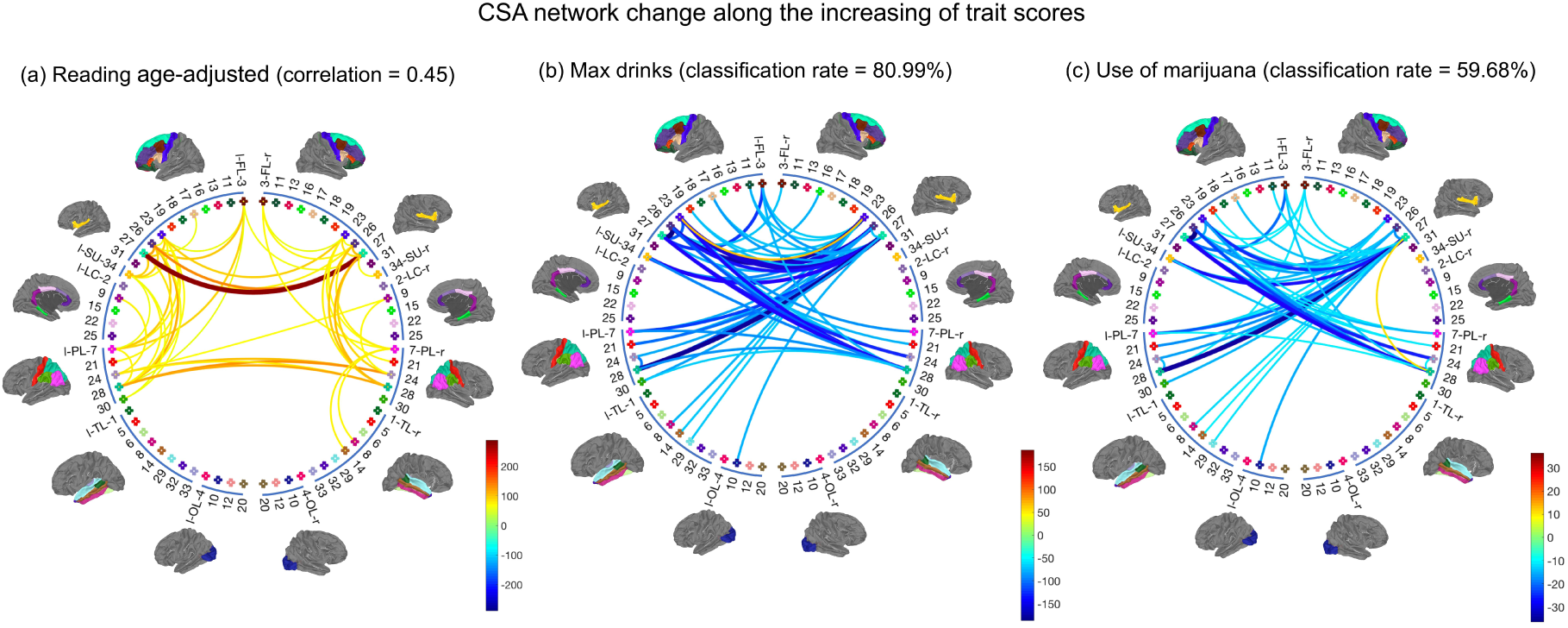
Top 50 pairs of brain regions in terms of their changes in CSA connectivity with increasing traits. (a) results for increasing language reading score. (b) results for max drinks; (c) results for marijuana use. We also display the correlation and binary classification rate obtained from CCA and LDA, providing measures of differences in brain connectivity networks between subjects with different trait scores.

From the networks presented in Figure 4, we extract their corresponding white matter tracts and display them in Figure 5. These white matter tracts are from two selected subjects with high and low trait scores. Among the connections shown in Figure 4, cross-hemisphere connections are particularly interesting. These connections are roughly classified into two types: (lFL, rFL) and (lPL, rPL). The first row of panel (a) in Figure 5 plots differences in these two types of connections between subjects with high (125.2) and low (67.27) reading scores. The streamlines corresponding to these connections are shown in the second row. The subject with a high reading score has richer and thicker structural connections in both (lFL, rFL) and (lPL, rPL); this pattern is common for subjects with similarly high reading scores. Figure 5 panel (b) shows similar results for selected light and binge drinkers; in this case differences between the subjects are detectable but subtle. More detailed results for the two pairs of subjects are displayed in the Supplementary Figure 5 and 6.

**Figure 5:**
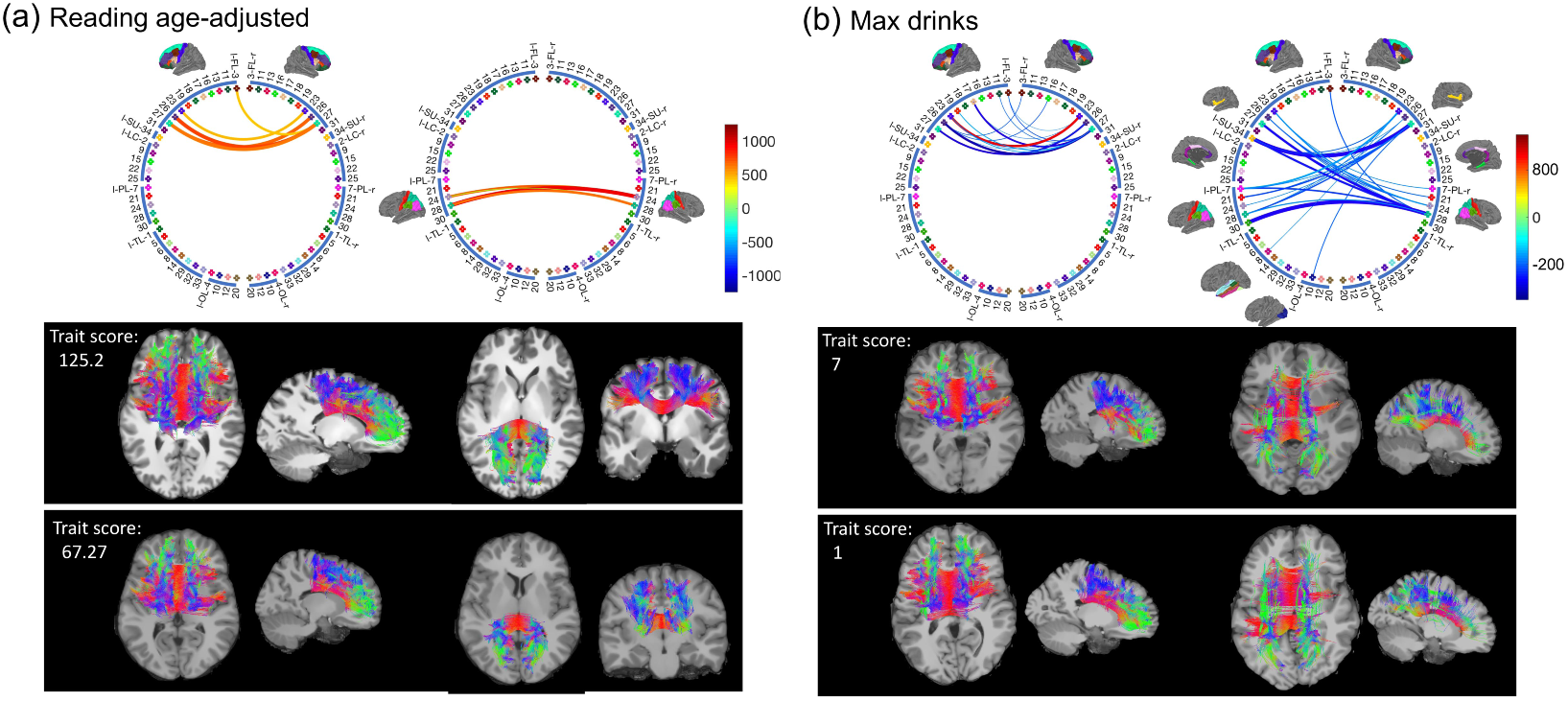
Network difference (based on CSA network) and the corresponding streamlines for selected subjects in the HCP data set. Among the 50 pairs of brain regions identified in 4, we focus on cross-hemisphere connections. Such connections are either (lFL, rFL) or (lPL, rPL). In (a) we show the differences in these two sets between subjects with high (125.2) and low (67.27) reading scores. The streamlines corresponding to these connections are shown in the second row. Similar results are plotted in (b) for two selected subjects with light and binge drinking.

## Discussion

Using state-of-the-art data science tools applied to data from the Human Connectome Project (HCP), we find that many different human traits are significantly associated with the brain structural connectome. Overall, and consistent with results in previous studies of functional connectomes ^[5,7]^, positive attributes tend to have an overall positive association with structural connectomes, while negative attributes have a negative relationship. Examples of positive traits include high language learning ability, fluid intelligence and motion ability; high levels of such variables tends to be indicative of stronger interconnections in the brain. Examples of negative traits include a high level of alcohol intake or the use of marijuana; such variables tend to be indicative of weaker interconnections.

Given inevitable errors in connectome reconstruction and in measuring human traits, such as alcohol intake, it is surprising how strong the statistical relationships are in our results. For example, we chose to highlight results for reading scores and alcohol intake as being particularly interesting. For reading scores using our data science methods, the correlation between the measured reading score and our predicted value based on an individual’s brain connectome was 0.45 (focusing on subjects with particularly low or high scores). In addition, and even more remarkably, the classification accuracy in attempting to distinguish between a light drinker and an individual with a history of binge drinking based only on their brain connectome was surprisingly high. (See Supplementary Figure 6 for more results on alcohol).

Code for implementing the data processing pipeline, TN-PCA method of dimensionality reduction, and the corresponding statistical methods for prediction and interpretation are all freely available with documentation on GitHub (to be posted on acceptance of this paper). These methods should be highly useful in further studies for carefully studying relationships between brain connectomes and individual traits. Careful follow-up studies are needed to better establish directions of causality. For example, do individuals with less connected brains have more of a tendency for substance abuse or does substance abuse cause a decrease in connectivity? The direction of this relationship has a fundamental impact on the clinical and public health implications of our results. Also of critical importance is the plasticity of the connectome; for example, if a binge drinker modifies their drinking behavior does the brain gradually return to a normal connectivity pattern over time? If an individual having a low reading score works hard to improve their score through coursework, tutoring and exercises, then does the brain connectivity also improve? Does this intervention have a direct causal effect on the connectome?

Given the increasing quality of data on structural connectomes, and in particular the sizable improvements in robustness and reproducibility, we are now at the point in which large, prospective studies can be conducted to answer some of the above important questions. The tools developed and used in this article should be helpful in analyzing data from such studies. On the methods side, it is important moving forward to continue to develop more informative and reproducible measures of connectivity between pairs of brain regions, and also to reduce sensitivity to the somewhat arbitrary number and choice of regions of interest. An additional important direction is linking structural and functional connectivity together in one analysis, which can potentially be accomplished via a minor modification of the proposed TN-PCA approach.

## Online Methods

### 0.1 Data sets

We focus on two data sets in this paper, which contain about 1, 133 dMRI scans for 1,067 subjects.

#### Human Connectome Project (HCP) Data set

The HCP aims to characterize human brain connectivity in about 1, 200 healthy adults and to enable detailed comparisons between brain circuits, behavior and genetics at the level of individual subjects. Customized scanners are used to produce high-quality and consistent data to measure brain connectivity. The latest release in 2017, containing various traits, structural MRI (sMRI) and diffusion MRI (dMRI) data for 1065 healthy adults, can be easily accessed through ConnectomeDB. The rich trait data, high-resolution dMRI and sMRI make it an ideal data set for studying relationships between connectomes and human traits.

A full dMRI session in HCP includes 6 runs (each approximately 10 minutes), representing 3 different gradient tables, with each table acquired once with right-to-left and left-to-right phase encoding polarities, respectively. Each gradient table includes approximately 90 diffusion weighting directions plus 6 *b*_0_ acquisitions interspersed throughout each run. Within each run, there are three shells of *b* = 1000, 2000, and 3000 s/mm^2^ interspersed with an approximately equal number of acquisitions on each shell. The directions are optimized so that every subset of the first *N* directions is also isotropic. The scan is done by using the Spin-echo EPI sequence on a 3 Tesla customized Connectome Scanner. See ^[37]^ for more details about the data acquisition of HPC. Such settings give the final acquired image with isotropic voxel size of 1.25 mm, and 270 diffusion weighted scans distributed equally over 3 shells.

In total, 175 covariates are extracted for each subject, including traits from domains of cognition, motor, substance use, psychiatric and life function, sensory, emotion, personality and health. Details about the extracted measures are presented in the Supplementary Material II.

#### Sherbrooke Test-Retest Data set

Different from the high-resolution HCP data set, this data set represents a clinical-like acquisition using a 1.5 Tesla SIEMENS Magnetom. There are 11 subjects with 3 acquisitions for each subject. A total of 33 acquisitions, from 11 healthy participants, were included. The diffusion space (q-space) was acquired along 64 uniformly distributed directions, using a b-value of *b* = 1000 s/mm^2^ and a single *b*_0_ (=0 s/mm^2^) image. The dMRI has a 2 mm isotropic resolution. An anatomical T1-weighted 1 × 1 × 1 mm^3^ MPRAGE (TR/TE 6.57/2.52 ms) image was also acquired. Diffusion data were up-sampled to 1 × 1 × 1 mm^3^ resolution using a trilinear interpolation and the T1-weighted image was registered on the up-sampled *b*_0_ image. Quality control by manual inspection was used to verify the registration.

### 0.2 Brain Connectome Extraction

#### HARDI tractography construction

A highly reproducible probabilistic tractography algorithm ^[9,18]^ is used to generate the whole-brain tractography data set of each subject for both data sets. The method borrows anatomical information from high-resolution T1-weighted imaging to reduce bias in reconstruction of tractography. Also the parameters are selected based on evaluation of various global connectivity metrics ^[18]^. In the generated tractography data, each streamline has a step size of 0.2 mm. On average, 10^5^ voxels were identified as the seeding region (white matter and gray matter interface region) for each individual in the HCP data set (with isotropic voxel size of 1.25 mm). For each seeding voxel, we initialized 16 streamlines to generate about 10^6^ streamlines for each subject. We have similar settings for the Sherbrooke test retest data set.

#### Network node definition

We use the popular Desikan-Killiany atlas to define the brain Regions Of Interest (ROIs) corresponding to the nodes in the structural connectivity network. The Desikan-Killiany parcellation has 68 cortical surface regions with 34 nodes in each hemisphere. Freesurfer software is used to perform brain registration and parcellation. Figure 1 column (a) illustrates the Desikan-Killiany parcellation and a reconstructed tractography data after subsampling.

#### Connectome tensor extraction

With the parcellation of an individual brain, we extract a set of weighted matrices to represent the brain’s structural connectome. To achieve this goal, for any two ROIs, one needs to first extract the streamlines connecting them. Alignment of the parcellation (on T1 image) and tractography data (dMRI image) is done using Advanced Normalization Tools (ANTs) ^[38]^. To extract streamlines connecting ROI pairs, several procedures are used to increase the reproducibility: (1) each gray matter ROI is dilated to include a small portion of white matter region, (2) streamlines connecting multiple ROIs are cut into pieces so that we can extract the correct and complete pathway and (3) apparent outlier streamlines are removed. Extensive experiments have illustrated that these procedures can significantly improve the reproducibility of the extracted connectomes, and readers can refer to ^[24]^ for more details.

To analyze the brain as a network, a scalar number is usually extracted to summarize each connection. For example, in the current literature ^[2,4,39]^, count is considered as a measure of the coupling strength between ROI pairs. However, the reliability of count is an issue due to the tractrography algorithms and the noise in the dMRI data ^[2,9]^. Instead of only using the count as the “connection strength”, we include multiple features of a connection to generate a tensor network for each brain. The tensor network has a dimension of *P* × *P* × *M*, where *M* represents the number of features and *P* represents the number of ROIs. Each of the M matrices is a weighted network and describes one aspect of the connection. As illustrated in the third column of Figure 1, the following features are extracted.

##### 1. Endpoint-related features

We consider the features generated from the end points of streamlines for each ROI pair. The first feature we extract is the number of end points, which is the same as the count of streamlines. Another feature we extracted is connected surface area (CSA), which is proposed in ^[24]^. To extract CSA, at each intersection between an ROI and a streamline, a small circle is drawn, and the total area covered by these circles is the CSA. A weighted version of CSA is calculated by dividing the total surface area of the two ROIs.

##### Diffusion-related features

Diffusion metrics, such as FA and generalized FA (GFA), characterize the water diffusivity at a particular location or voxel. We extract these diffusion metrics along white matter streamlines for each ROI pair. For example, the mean of FA, the max FA, the mean of MD and the max MD are calculated as weighted networks.

##### Geometry-related features

The average length, shape, and cluster configuration ^[24]^ characterize the geometric information of streamlines for each ROI pair. The average length of streamlines reflects the intrinsic spatial distance between two regions. The number of streamline clusters is more robust to some confounding effects in the tractography reconstruction, e.g. the seeding strategy. In addition, the deviations of streamline shapes from a mean streamline are interesting features and we extract them as the shape features (see ^[24]^ for more details).

### 0.3 Tensor-Network Principal Components Analysis

In this section, we develop a Tensor Network PCA (TN-PCA) method to embed brain networks into low dimensional vector spaces. Just as matrix decompositions are used for PCA, tensor decompositions have been widely used for PCA with tensor data ^[40]^. Here, we develop a new tensor decomposition that is specifically appropriate for our tensor network data.

To begin, we need some notation which is largely adapted from ^[40]^. Let ***χ*** ∈ ℝ ^*I*_1_×*I*_2_×…×*I*_*N*_^ be an *N-mode* tensor, and we denote matrices as **X**, vectors as x and scalars as *x.* The *outer product* is denoted by ○: x ○ y = xy^T^. The *scalar product* of two tensors ***A, B*** ∈ ℝ ^*I*_1_×*I*_2_×…×*I*_*N*_^is defined as 〈***A, B*** 〉 = Σ_*i*_1__ Σ_*i*_2__ … Σ_*i*_*N*__ *a*_*i*_1_*i*_2_…*i*_*N*__*b*_*i*_1_*i*_2_…*i*_*N*__. The *Frobenius norm* of a tensor ***χ*** is 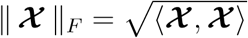. The *n-mode multiplication* of tensor ***χ*** ∈ ℝ^*I*_1_×*I*_2_×…×*I*_*N*_^ with a matrix **A** ∈ ℝ^*J*_*n*_×*I*_*n*_^, denoted by ***χ*** ×_*n*_**A**, gives a tensor in ℝ^*I*_1_×…*I*_*n*–1_×*J*_*n*_×*I*_*n*+1_…×*I*_*N*_^, where each element is the product of mode-*n* fiber of *χ* multiplied by **A**. When **A** degenerates to a vector a, i.e. a ∈ ℝ^*I*_*n*_^, we have ***χ***×_*n*_a ∈ ℝ^*I*_1_×…*I*_*n*–1_×*I*_*n*+1_…×*I*_*N*_^, which is an *N* – 1-mode tensor. For simplicity, we will only consider three-mode tensors here, but all of the methods can be easily extend to higher-order tensors. We will work with a tensor network, ***χ*** ∈ ℝ^*P*×*P*×*N*^, which is a concatenation of network adjacency matrices **A**_*i*_ ∈ ℝ^*P*×*P*^ for *i =* 1,…, *N*, where *P* represents the number of nodes and *N* represents the number of subjects (or features we extracted from each connection). We say that our tensor network is *semi-symmetric* as every frontal slice, ***σ***_:,:, *n*_, is a symmetric matrix: ***χ***_*i,j,n*_ = ***χ***_*j,i,n*_∀ *i, j, n*.

Given the special structure of our semi-symmetric tensor network, existing tensor decompositions are not ideal for conducting tensor PCA. Consider an extension of the popular *Tucker* model ^[41]^ which forces the first two Tucker factors to be equal to account for the semi-symmetric structure:

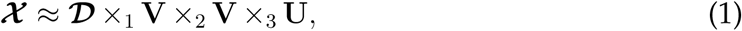

where V_*P*×*K*_*V*__ and U_*N*×*K*_*U*__ are orthogonal matrices that form the Tucker factors and ***D***_*K*_*V*_×*K*_*V*_×*K*_*U*__ is the Tucker core; under certain restrictions on the Tucker core, this model will result in a semi-symmetric tensor. Interestingly, when standard algorithms for estimating Tucker models (e.g. Higher-Order SVD and Higher-Order Orthogonal Iteration, HOSVD and HOOI, respectively ^[41-43]^) are applied to semi-symmetric tensors, they result in tensor factorizations that follow model (1); this fact can be easily verified and is also discussed in ^[42]^. Despite the ease of implementation of Tucker models for semi-symmetric tensors, these approaches are not ideal for studying and embedding brain networks. The semi-symmetric Tucker core makes it difficult to directly interpret the effects and prevalence of tensor network components (eigenvectors associated with V) across the population. Further, the assumption that the population factors, U, be orthogonal is likely overly restrictive and limits the Tucker model’s ability to fit data well in our application.

Because of this, we consider another popular tensor decomposition model: the CP decomposition, which models a tensor as a sum of rank one tensors: 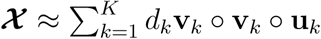 ^[44,45]^. As with the Tucker model, it is clear that the first two factors must be equivalent to yield a CP model appropriate for semi-symmetric tensors:

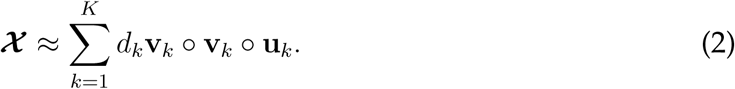

Here, v_*k*_ and u_*k*_ are *P* and *N* vectors, respectively, that form the *k^th^* CP factor and *d*_*k*_ is the *k*^*th*^ positive CP scaling parameter. It is clear that (2) needs no further restrictions to yield a semi-symmetric tensor. Yet, this model may not be ideally suited to modeling our population of brain connectomes. If there are no restrictions on the CP factors V as is typical in CP models, then the columns of V could be highly correlated and fail to span the eigen-space of the series of brain networks. Hence, we propose to add an additional orthogonality constraint on the CP factors V, but leave U unconstrained. Note that this form of orthogonality in one factor but not the other is distinct from the various forms of orthogonal tensor decompositions proposed in the literature ^[46]^.

We estimate our CP model for semi-symmetric tensors by solving the following least squares problem:

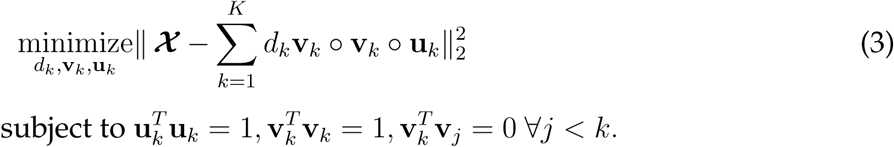

As with the typical CP problem, this is non-convex but is instead bi-convex in v and u. The most common optimization strategy employed is block coordinate descent which alternates solving a least squares problem for all the *K* factors, for V with U fixed and for U with V fixed ^[43]^. For our problem with the additional orthogonality constraints, this approach is computationally prohibitive. Instead, we propose to use a greedy one-at-a-time strategy that sequentially solves a rank-one problem, a strategy sometimes called the tensor power method ^[47,48]^. The single-factor CP method can be formulated as

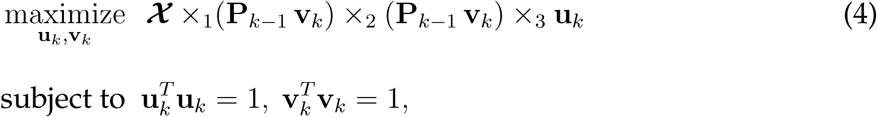

where 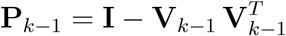 is the projection matrix with V_*k*–1_ = [v_1_,… v_*k*–1_] denoting the previously estimated factors. (4) uses a Gram-Schmidt scheme to impose orthogonality on v_*k*_ via the projection matrix *P*_*k*–1_; it is easy to verify that (4) is equivalent to (3) ^[48]^.

To solve (4), we employ a block coordinate descent scheme by iteratively optimizing with respect to u and then v; each coordinate-wise update has an analytical solution:

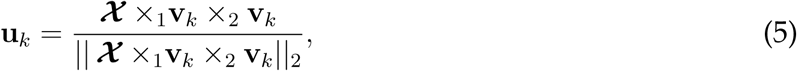

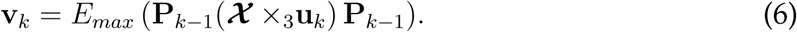

Here, *E*_*max*_(A)refers to the maximum eigenvalue of matrix A. We can show that this scheme converges to a local optimum of (4). Putting together these pieces, we present our tensor power algorithm for solving (3) in Algorithm 1. After we greedily estimate a rank-one factor, we use subtraction deflation. Overall, this algorithm scales well computationally compared to the Tucker model which requires computing multiple SVDs of potentially large matricized tensors.

#### Algorithm 1 Tensor power method for semi-symmetric CP decomposition (TN-PCA Algorithm)

Let ***χ*** be a three-way tensor concatenating *M* brain networks. The tensor power method for semi-symmetric CP decomposition of ***χ*** is given as:

1. Let 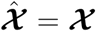.
2. For *k* = 1, …, *K*, do
  a. (a) Find the semi-symmetric single-factor CP decomposition for 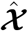. Initialize v_*k*_, u_*k*_, and iteratively update v_*k*_, u_*k*_ until convergence:

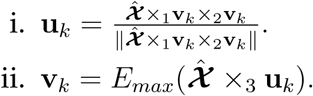
  b. CP Scaling: 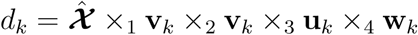
  c. Projection: 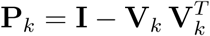
  d. Deflation: 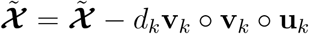

When applied to tensor brain networks, our new semi-symmetric tensor decomposition results in a method for tensor-network PCA (TN-PCA). Specifically, each u_*k*_ denotes the *subject-mode* and each 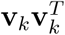 denotes the rank-one *network mode.* The subject-modes give the low dimensional vector embeddings of the brain networks for each subject; we use these to associate structural connectomes with traits. We call the weighted sum of network modes, 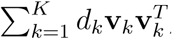, the principal brain network, which gives a one network summary that captures the most variation in the structural connectomes across all subjects. We empirically validate our proposed CP decomposition for semi-symmetric tensors and compare this to the existing HOSVD and HOOI approaches through simulations and an analysis of the Sherbrooke Test-Retest data set in the supplementary materials. Overall, the results indicate that our method yields a more flexible model for tensor brain networks that out-performs existing methods.

### 0.4 Relating Connectomes to Traits

Let *χ* be a three-way tensor with dimension of *P* × *P* × *N*, representing the stacking of *P* × *P* weighted networks from *N* subjects. The TN-PCA 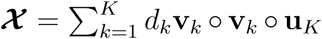 embeds the weighted networks into vectors U_*K*_ = [u_1_, …, u_*k*_], where each row represents one embedded coordinate for a weighted network in ℝ^*K*^. Since we have restricted {v_*k*_} to be orthogonal to each other, we have (v_*k*_ ○ v_*k*_) * (v_*k*’_ ○ v_*k*’_) = 0 for *k* ≠ *k*’. Therefore, another way to view TN-PCA is that {v_*k*_ ○ v_*k*_} are basis networks and u_*k*_(*i*) for *i* = 1, …, *N* are the corresponding normalized coefficients (for *i*th subject) and the scale is absorbed by *d*_*k*_.

To relate the networks to various traits, we rely on the embedded vectors U_*K*_. There are many advantages of using the vector U_*K*_(*i*,:) to represent the *i*th subject’s brain connectivity, e.g., (i) we recover the network from U_*k*_(*i*,:) through 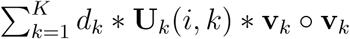; (ii) the low-dimensional vector representations bring us flexibility to utilize various existing statistical tools to study the relationship between the brain networks and human traits; and (iii) the TN-PCA is flexible and can be easily extended to deal with high-order tensors, e.g. a four-mode tensor, by simply include another vector in the outer product.

#### Hypothesis testing of distribution difference

For each trait, we sort the 1065 HCP subjects according to their scores, and extract two groups of subjects: 100 subjects with the highest trait scores and 100 subjects with the lowest scores. For discrete traits, it is sometimes not possible to identify exactly 100 subjects; in such cases, we randomly select subjects on the boundary as needed. We compare the embedded vectors of networks (rows of U_*K*_) from the two groups and test the null hypothesis that the two samples are from the same distribution against the alternative that they are from different distributions. We use the Maximum Mean Discrepancy (MMD) ^[31]^ to perform hypothesis tests. FDA is controlled using ^[32]^.

#### Relating brain connectomes with traits

We are interested in studying relationships between traits and brain connectomes for all HCP subjects. Particularly, we are interested in two scenarios. First, we want to see if brain connectomes can be used to predict various traits. Second, for those traits that can be predicted by brain connectomes, we would like to identify how the connectome changes with trait values by flagging the subset of connections with the largest differences.

Let **Y** = [*y*_1_, …, *y*_*N*_]^*T*^ be a *N* × 1 vector, which contains trait scores for *N* subjects, and let U_*K*_ be an *N* × *K* matrix containing the embedded networks in ℝ^*K*^ for *N* subjects. Our first analysis focuses on predicting *y*_*i*_ using brain PC scores and other demographic covariates such as age and gender. Subjects in the HCP are randomly allocated into a training data set (66% of the subjects), a validation data set (17% of the subjects) and a test data set (17% of the subjects). We then train various machine learning methods (e.g., simple linear/logistic regression, random forests, support vector machines and XGboost) to predict the trait scores. The prediction accuracy is evaluated using the root-mean-square error (RMSE) for both continuous and ordinal traits and the classification accuracy for categorical traits. To evaluate whether brain connectomes are important in predicting traits, we compare with a reduced model, where only some demographic covariates are used as predictors. The model containing brain connectomes is referred as the full model, and the model without brain connectomes is referred to as the baseline model.

For each trait, we define a measure *ρ* to evaluate the importance of brain connectomes. For trait *p*, let 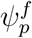 denote the RMSE or (1- classification accuracy) of the full model, and 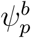 for the baseline model. The measure *ρ* for trait *p* is calculated as 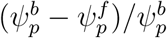. The best model (including the tuning parameters in each machine learning method) is selected based on the validation data set, and then is applied to the test data set for performance evaluation. We then report *ρ* for different combinations of networks and traits. Figure 4 in the Supplementary Material I shows *ρ* for both validation and test data sets for each trait. The results were based on the average of 50 runs. The Supplementary Material II summarizes details of the 175 traits.

For each trait, we identify a subset of edges that change significantly with the trait. Depending on a trait’s type, we apply different methods to identify the subset of edges. For continuous traits we use canonical correlation analysis (CCA) ^[33]^ and for categorical traits we use linear discriminant analysis (LDA) ^[49]^. For a continuous trait, the problem of finding a subset of edges that are highly correlated with the trait using the CCA method is equivalent to finding a direction w in *ρ*^*K*^ (in the network embedding space) such that the correlation between the trait scores {*y*_*i*_} and the projection scores {*u*_*proj*_(*i*) = 〈U_*K*_(*i*,:), w〉} are maximized, i.e.
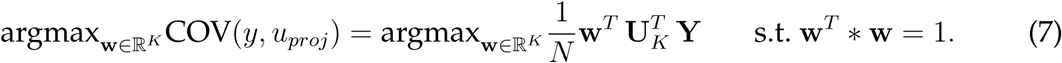

If we assume that both Y and rows of U_*K*_ are centered, the unit vector w obtained in (7) describes changes of in networks in the embedding space with increases of the trait score. To obtain the corresponding change in each edge in the network form ∆_*net*_, we let 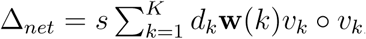, where *s* ∈ ℝ^+^ is a scalar that we will define later. Confounding influence of age and gender can be regressed out. For a categorical trait, e.g., *y*_*i*_ ∈ {0,1}, we use LDA to identify edge changes from low to high scores. The idea is to find a w in ℝ^*K*^ that best separates the two classes of networks. Let *μ*_0_ and *μ*_1_ be the means, and Σ_0_ and Σ_1_ be the covariances of the embedded networks. The separation of the two groups is defined in the embedding space in the following way:

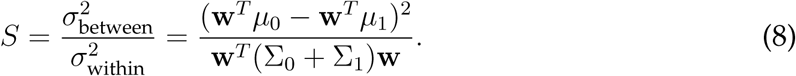

It is clear that *S* achieves the maximum when w ∝ (Σ_0_ + Σ_1_)^−1^(*μ*_1_ - *μ*_0_).

In both CCA and LDA methods, since we only identify a unit vector w, which only reflects the global trend of the network changes with increasing traits and there is no scale information included in w. This setting makes the comparison of the network changes across different traits hard. Hence, we want to define a proper scale *s* for each trait such that ∆_*net*_ reflects the magnitude of network changes with increases in that particular trait.

Since in this paper, the w’s are always inferred from two groups of subjects (with high and low trait scores even for continuous traits), we binarize the subjects and use their mean differences in the network embedding space to define *s*. To be more specific, for a selected trait, letting ū_0_, ū_1_ ∈ ℝ^*K*^ be the mean embedding vectors for subjects with low and high trait scores respectively, we define *s* = ‖ū_0_ - ū_1_‖. In Figure 4 of the main paper, we plot the *∆*_*net*_ for each trait.

## Acknowledgements

Data were provided in part by the Human Connectome Project, WU-Minn Consortium (Principal Investigators: David Van Essen and Kamil Ugurbil; 1U54MH091657). We also thank Maxime Descoteaux, Kevin Whittingstall and the Sherbrooke Molecular Imaging Center for the acquisition and sharing of the test-retest data.

